# ADAM17 targeting by human cytomegalovirus remodels the cell surface proteome to simultaneously regulate multiple immune pathways

**DOI:** 10.1101/2023.03.16.532955

**Authors:** Anzelika Rubina, Mihil Patel, Katie Nightingale, Martin Potts, Ceri A. Fielding, Simon Kollnberger, Betty Lau, Kristin Ladell, Kelly L. Miners, Jenna Nichols, Luis Nobre, Dawn Roberts, Terrence M. Trinca, Jason P. Twohig, Virginia-Maria Vlahava, Andrew J. Davison, David A. Price, Peter Tomasec, Gavin W.G. Wilkinson, Michael P. Weekes, Richard J. Stanton, Eddie C.Y. Wang

**Author notes:** Deceased (Peter Tomasec). These authors contributed equally to this study (Anzelika Rubina, Mihil Patel, Richard J. Stanton, Eddie C.Y. Wang).

## Abstract

Human cytomegalovirus (HCMV) is a major human pathogen whose life-long persistence is enabled by its remarkable capacity to systematically subvert host immune defences. In exploring the finding that HCMV infection upregulates tumor necrosis factor receptor 2 (TNFR2), a ligand for the pro-inflammatory anti-viral cytokine TNFa, we discovered the underlying mechanism was due to targeting of the protease, A Disintegrin And Metalloproteinase 17 (ADAM17). ADAM17 is the prototype ‘sheddase’, a family of proteases that cleaves other membrane-bound proteins to release biologically active ectodomains into the supernatant. HCMV impaired ADAM17 surface expression through the action of two virally-encoded proteins in its *U_L_*/*b*’ region, UL148 and UL148D. Proteomic plasma membrane profiling of cells infected with a HCMV double deletion mutant for UL148 and UL148D with restored ADAM17 expression, combined with ADAM17 functional blockade, showed that HCMV stabilized the surface expression of 114 proteins (p<0.05) in an ADAM17-dependent fashion. These included known substrates of ADAM17 with established immunological functions such as TNFR2 and Jagged1, but also numerous novel host and viral targets, such as Nectin1, UL8 and UL144. Regulation of TNFα-induced cytokine responses and NK inhibition during HCMV infection were dependent on this impairment of ADAM17. We therefore identify a viral immunoregulatory mechanism in which targeting a single sheddase enables broad regulation of multiple critical surface receptors, revealing a paradigm for viral-encoded immunomodulation.

**Significance statement:** Human cytomegalovirus (HCMV) is an important pathogen, being the commonest infectious cause of brain damage to babies and the primary reason for hospital readmissions in transplant recipients. Even though HCMV induces the strongest immune responses by any human pathogen, it evades host defences and persists for life. This study describes a novel immunoregulatory strategy through which HCMV modulates multiple immune pathways simultaneously, by targeting a single host protein. HCMV UL148 and UL148D impair the maturation of the sheddase, A Disintegrin And Metalloproteinase 17, profoundly altering surface expression of numerous immunoregulatory proteins. This is the first description of viral genes targeting this pathway. Our findings may be relevant for future viral therapies and understanding the impact of HCMV in developmental biology.

## INTRODUCTION

Human cytomegalovirus (HCMV) is the leading infectious cause of congenital birth defects and is responsible for morbidity and mortality in immunocompromised individuals, in particular transplant recipients (1–3). HCMV persists lifelong after primary infection in the face of potent innate, humoral and cell-mediated immunity. A large proportion of HCMV’s substantial gene content is dedicated to manipulating host immune defences. HCMV immunevasins disrupt natural killer (NK) and T cell responses by impairing antigen presentation, downregulating expression of activating ligands, upregulating ligands for inhibitory receptors, and altering the functionality of host proteins to manipulate antiviral signalling and immune pathways. Ultimately, HCMV immunevasins have a profound impact on the capacity of both infected cells and immunological effector cells to respond to infection (4–7). They continue to be the subject of intense research with the hope that the understanding gained will aid in generating new treatments against HCMV.

Tumor Necrosis Factor alpha (TNFα) is a primary effector cytokine produced by NK and T cells. The TNFa pathway plays a key role in inflammation and anti-viral immunity, as evidenced by the increased broad viral reactivation (8), but also HCMV driven inflammatory disorders (retinitis, hepatitis, ileitis, colitis), observed in patients undergoing anti-TNF treatment (9–12). The effects of TNFa are achieved following binding with two receptors, TNFR1 and TNFR2. The outcome of TNFR1 signalling is context dependent, as TNFR1 contains intracellular motifs capable of recruiting both downstream promoters of apoptosis (TNFR1-associated death domain, Fas-associated death domain) and also survival leading to pro-inflammatory immune responses (TRAF2, NF-κB) (13, 14). Both these outcomes should apparently counter viral infection, however HCMV exploits this pathway by encoding at least 5 genes that block apoptosis (15–18), and upregulating surface expression of TNFR1 using the viral protein UL138, which then aids TNFα-mediated HCMV reactivation (19). The role of TNFR2 during the HCMV life cycle is more obscure yet is as important to understand considering its inability to induce apoptosis, but capacity to trigger the NF-κB pathway (20, 21). In contrast to TNFR1 which is found constitutively on most nucleated cells, TNFR2 expression is inducible and primarily limited to immune cells (22, 23). Dynamic changes in the levels of TNFR2 and subsequent responses to TNFα therefore have the potential to significantly modulate immune responses (24) as well as affect the HCMV life cycle.

The initial aim of this study was to investigate the regulation of TNFR2 during HCMV infection, however this led to the discovery that its altered levels were an indirect consequence of the targeting of A Disintegrin And Metalloproteinase 17 (ADAM17). ADAM17 is an ectodomain shedding protease with established biological importance in both mice and humans. There are over one hundred known ADAM17 substrates including cell adhesion molecules (e.g. L-selectin), immunoregulatory cytokines including TNFα, and cytokine receptors such as TNFR1 and TNFR2 (25, 26). We thus describe the identification of two viral genes responsible for down-regulating ADAM17, detailing the profound effect this has on the cell-surface proteome, and defining some of the important immunological consequences of this in the context of HCMV infection.

## RESULTS

### Surface upregulation of TNFR2 by HCMV is dependent on UL148 and UL148D

We began our investigations by studying cell surface expression of TNFR2 during HCMV infection. The high passage laboratory strains AD169 and Towne induced a modest increase in TNFR2 on the cell surface (27), yet a much more substantial upregulation was observed with Toledo and the low passage strain Merlin (Fig. 1A). Strains AD169 and Towne have suffered ~15kb and ~13kb deletions, respectively, at one end of the UL segment, termed the *U_L_*/*b*’ region (28). Although *U_L_*/*b*’ has suffered an inversion in Toledo, it remains otherwise intact. We therefore hypothesised that function(s) encoded by *U_L_*/*b*’ modulated the expression of TNFR2. To determine the specific HCMV genes involved, we performed a loss of function screen by assessing the levels of surface TNFR2 in cells infected with a library of HCMV strain Merlin single gene deletion mutants spanning the entire *U_L_*/*b*’ region (Fig. 1B). While all the deletion mutants downregulated surface expression of HLA-I (an indicator of HCMV infection), only ΔUL148 and ΔUL148D showed reduced levels of TNFR2 compared to the Merlin-infected control (Fig. 1B, C). A time-course analysis indicated that differencesin TNFR2 expression could be detected as early as 24h pi and continued to increase through to 72 hrs pi (Fig. 1D).

**Figure 1.**
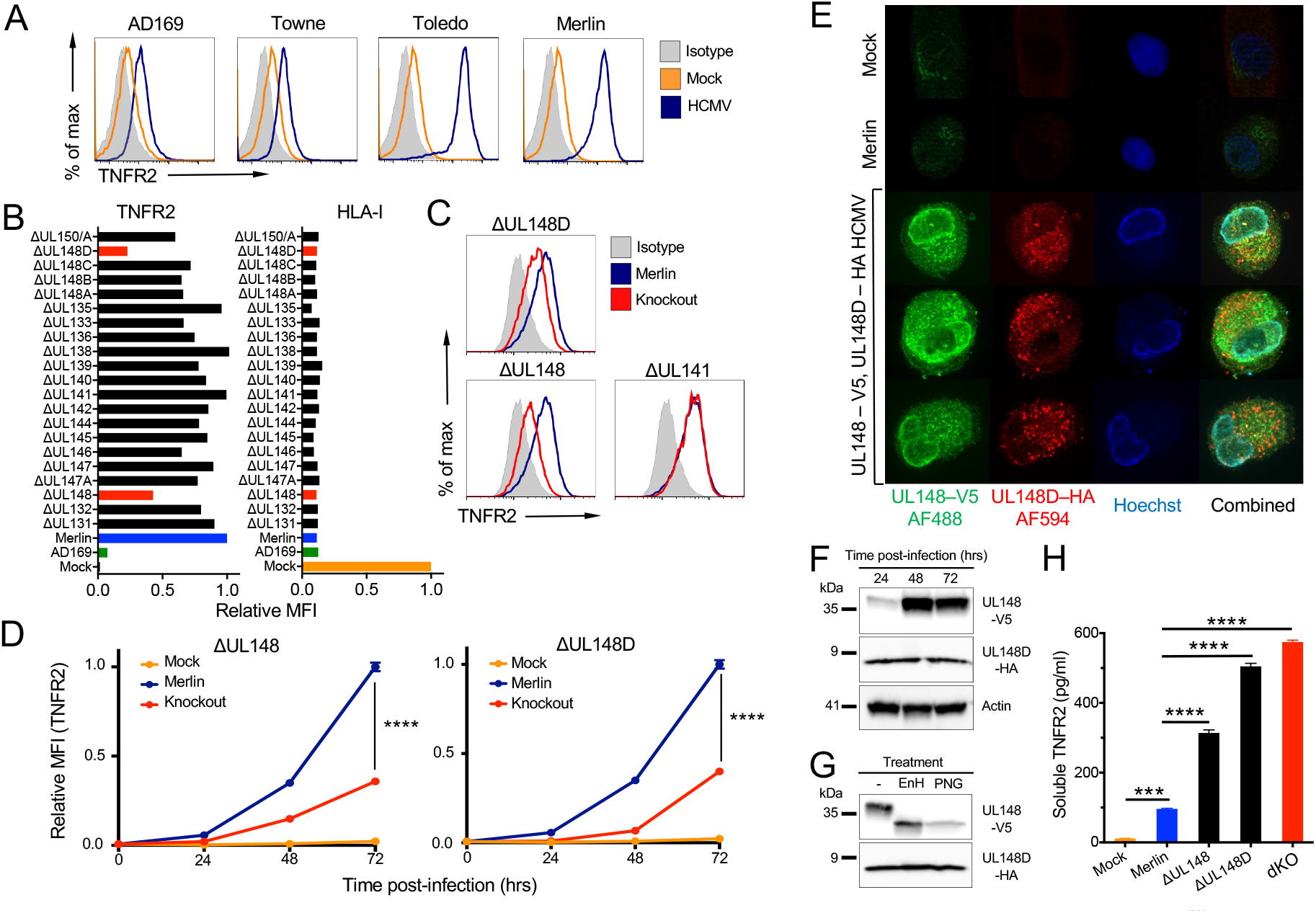
Legend: HCMV genes UL148 and UL148D upregulate surface expression of TNFR2. (A) Flow cytometric histogram overlays showing surface TNFR2 expression of HF-TERT cells, mock-infected or infected with the indicated HCMV strains (MOI=10, 72 h pi). (B) Relative surface expression of TNFR2 and HLA-I on HF-TERT cells infected with a library of HCMV strain Merlin deletion mutants. Median fluorescence intensity (MFI) values from flow cytometric histograms are shown relative to Merlin-infected cells (set to 1) for TNFR2 or mock-infected cells (set to 1) for HLA-I. (C) Flow cytometric histogram overlays showing surface TNFR2 expression of HF-TERT cells infected with the indicated HCMV strain Merlin deletion mutants (MOI = 5, 24 h pi). (D) Relative surface expression of TNFR2 with time of HF-TERT cells, mock-infected or infected with the indicated HCMV strains. MFI values from flow cytometric histograms are shown relative to Merlin-infected cells (set to 1) at 72 h pi. Data are shown as mean ± SEM of triplicate infections. Two-way ANOVA (with Dunnett’s T3 multiple comparison post-hoc test showed significance at \*\*\*\**p*<0.0001. (E) Fluorescence microscopy for UL148 and UL148D in HF-TERT cells, mock-infected or infected with HCMV strain Merlin or HCMV Merlin UL148-V5/UL148D-HA. At 48 h pi, cells were fixed, permeabilized, and stained with anti-V5 and anti-HA antibodies, then Alexa Fluor 488-conjugated anti-rabbit IgG and Alexa Fluor 594-conjugated anti-mouse IgG, before counterstaining with Hoechst. (F, G) Total protein expression of UL148 and UL148D in HF-TERT cells infected with HCMV Merlin UL148-V5/UL148D-HA. Whole-cell lysates were analyzed by immunoblotting at the indicated time points pi (F) or at 72 h pi after digestion of the lysates with EndoH or PNGaseF (G). (H) HF-TERT cells were mock-infected or infected with HCMV strain Merlin or the indicated deletion mutants, and levels of soluble TNFR2 (sTNFR2) in the culture medium at 72 h pi measured by ELISA.

To characterise UL148 and UL148D expression during infection, we engineered C-terminal V5 and HA tags onto UL148 and UL148D, respectively, within a single recombinant HCMV. Immunofluorescence indicated UL148 and UL148D were both excluded from the nucleus but did not substantially co-localize within the infected cell, with UL148D showing more vesicular staining (Fig. 1E). The differences in localization were mirrored by differences in patterns of protein expression as determined by Western blot analysis. UL148 expression was low at 24 hrs pi but increased as infection progressed, reaching high levels by 48 hrs pi (Fig. 1F). UL148 ran as a glycosylated ~35 kDa protein that was Endo-H sensitive, consistent with residence in the ER (Fig. 1G). In contrast, UL148D expression had plateaued by 24 hrs pi (Fig. 1F), running around ~7 kDa with no evidence of glycosylation (Fig. 1G).

Cell surface expression of TNFR2 was upregulated more by HCMV ΔUL148 and ΔUL148D than by strains missing the U_L_/b’ region, although none of these viruses attained the levels induced by wild-type HCMV Merlin (Fig. 1D). We therefore engineered a ΔUL148/ΔUL148D double knockout (dKO) HCMV mutant, infection with which resulted in TNFR2 levels comparable to those detected using strain AD169 (Supplemental Fig. 1A, B). UL148 and UL148D thus both contributed to the upregulation observed with wild-type virus. The abundance of TNFR2 detected in whole cell lysates was nevertheless similar whether HCMV encoded UL148 and UL148D, or not (Supplemental Fig. 1C). This observed upregulation of TNFR2 by HCMV Merlin was therefore limited to the cell surface. Since TNFR2 may be shed from cells, soluble TNFR2 (sTNFR2) levels in infected cell supernatants were also assessed. While HCMV infection increased sTNFR2 levels, concentrations were enhanced in the absence of UL148 and/or UL148D (Fig. 1H). These collective findings implied that UL148 and UL148D may promote high levels of surface TNFR2 expression by impeding its shedding.

### UL148 and UL148D target ADAM17

ADAM17 controls the surface expression of select proteins via proteolytic cleavage/release of their ectodomains, including TNFR1 and TNFR2 (25, 26). When we compared the cell surface proteomes of human fibroblasts infected with HCMV ΔUL148, HCMV ΔUL148D and the parental virus Merlin, ADAM17 surface expression was increased in each individual HCMV gene knockout (Fig. 2A). Consistent with this, flow cytometry demonstrated complete abolishment of surface ADAM17 on wild-type HCMV Merlin-, but its recovery on dKO- and AD169-, infected cells (Fig. 2B, C). Moreover, ectopic expression of either UL148 or UL148D using adenovirus vectors (rAds) resulted in downregulation of surface ADAM17, with a more marked reduction when both genes were expressed together (Fig. 2D). UL148 and UL148D therefore independently downregulated ADAM17 from the cell surface but could act in concert to achieve a greater effect (Fig. 2D, E).

**Figure 2.**
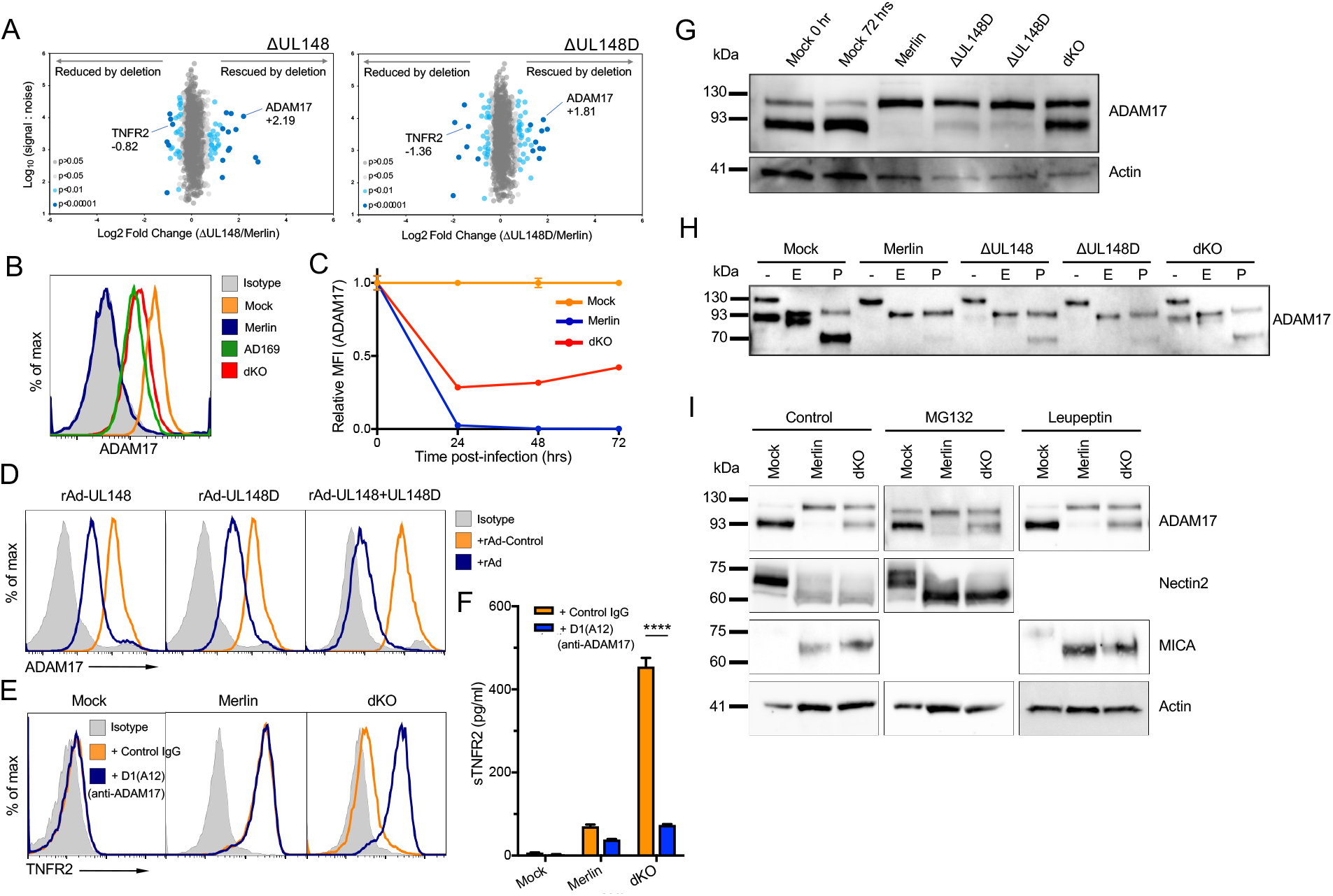
Legend: UL148 and UL148D upregulate surface TNFR2 by impairing the maturation of ADAM17. (A) Scatterplot of cell-surface proteins modulated by UL148 and UL148D. Data were generated using PMP. Fold change was calculated for each protein by comparing the signal:noise (S:N) value from each sample infected with a deletion virus to the S:N value for the same protein from the sample infected with HCMV strain Merlin. Benjamini-Hochberg-corrected significance B was used to estimate p-values. This metric calculates the probability of obtaining a log-fold change of at least a given magnitude under the null hypothesis that the distribution of log-ratios has normal upper and lower tails. Modifications allowed the spread of upregulated and downregulated values to be different, and values were calculated for consecutive protein subsets obtained by sequential S:N binning, because the spread of fold change ratios for proteins quantified by peptides with high S:N values is naturally smaller than the spread of fold change ratios for less well quantified proteins with lower total S:N values (79). (B, C) Surface expression of ADAM17 on HF-TERT cells, mock-infected or infected with the indicated HCMV strains as shown by flow cytometric overlay histogram at 72 h pi (B), and relative to MFI of mock-infected cells at 72 h pi (set to 1) at the indicated time points (C). Data are shown as mean ± SEM of triplicate infections. (D) Flow cytometric overlay histograms showing ADAM17 expression on HF-CAR cells infected with a control vector (RAd-Control) or RAds encoding UL148, UL148D or a combination of both (MOI=10, 72 h pi). (E) Flow cytometric overlay histograms showing TNFR2 expression on HF-TERT cells, mock-infected or infected with indicated HCMV strains for 48 h before addition of anti-ADAM17 antibody D1(A12) or human IgG for additional 24 h. (F) Soluble TNFR2 levels in media of cultures as in (E). Data are shown as mean + SEM of triplicate samples. Two-way ANOVA with a Bonferroni post-test showed significance at \*\*\*\**p*<0.0001. (G) Whole-cell protein levels of ADAM17 visualized by immunoblotting using lysates from HF-TERTs infected with the indicated HCMV strains for 72 h. Actin was used as a loading control. (H) Western blots for ADAM17 of whole-cell lysates generated as in (G) but treated with EndoH or PNGaseF. (I) Western blots of whole-cell lysates from HF-TERT cells, mock-infected or infected with HCMV strain Merlin or the dKO mutant for 72 h, with treatment of proteasomal (MG132) or lysosomal (leupeptin) protein degradation inhibitors for the last 12 h. Immunoblotting for ADAM17, nectin2 (positive control for MG132 treatment), MICA (positive control for leupeptin treatment), and actin as a loading control.

To further test whether cell surface upregulation of TNFR2 was indeed caused by the abrogation of ADAM17 function, we used an antibody, D1(A12), that specifically inhibits the shedding activity of ADAM17 (29). D1(A12) treatment of cells infected with the dKO mutant, resulted in the recovery of surface TNFR2 comparable to levels observed following infection with the parental Merlin virus (Fig. 2E). Application of D1(A12) also reduced levels of sTNFR2 in the supernatants from cells infected with the dKO mutant, compared to Merlin (Fig. 2F). Alterations in both cell surface and soluble TNFR2 induced by HCMV infection were therefore consequent to the targeting of ADAM17 by UL148 and UL148D.

### Intracellular retention of ADAM17 during HCMV infection

ADAM17 is expressed as a precursor that is itself proteolytically cleaved (30). Immunoblotting identified two differentially migrating forms of ADAM17 in mock-infected cells (Fig. 2G). The slower migrating species exhibited partial EndoH sensitivity, consistent with it being an immature, intracellular precursor (Fig 2G, H). The faster migrating form was EndoH resistant, which is consistent with it being the cleaved, fully glycosylated mature functional form of ADAM17. This mature form was absent in Merlin-infected cells, indicating that ADAM17 processing had been impaired during wild-type HCMV infection. Infection with either ΔUL148 or ΔUL148D was associated with partial recovery of the mature form, while infection with dKO HCMV resulted in its full recovery, detected as a strong low mwt signal (Fig. 2G). Thus, UL148 and UL148D both impair the maturation of ADAM17 from its intracellular precursor.

UL148 has previously been shown to interact with proteins in the ERAD pathway (31, 32). We therefore investigated the role of proteasomal degradation in UL148/UL148D-driven loss of mature ADAM17. Treatment with the proteasome inhibitor MG132 recovered expression of nectin2 in Merlin-infected cells as described previously (33), but did not recover mature ADAM17 (Fig. 2I). ADAM17 is also passed down the lysosomal pathway of destruction following cellular activation (34), so we treated Merlin-infected cells with the lysosomal inhibitor leupeptin. This recovered expression of MHC class I chain-related protein A (MICA), previously shown to be targeted to the lysosomal pathway by HCMV (35), but there was no recovery of mature ADAM17 (Fig. 2I). Taken together, these data suggest that the lack of surface ADAM17 expression during wild-type HCMV infection is not caused by degradation of its mature form, but retention and absence of processing of its immature form inside the cell.

### Impact of UL148 and UL148D on the surface proteome

The intracellular retention of ADAM17 mediated by UL148 and UL148D may be expected to affect the expression of additional viral and cellular protein during infection. Plasma membrane proteomics (PMP) was therefore performed on cells infected with HCMV ΔUL148, HCMV ΔUL148D, the dKO virus and the parental Merlin strain. In addition, dKO-infected cells were treated with D1(A12) (Fig. 3A). Cell surface proteins impacted by the loss of UL148/UL148D and rescued by treatment with D1(A12) were defined as targets manipulated specifically by UL148/UL148D-mediated targeting of ADAM17 (Fig. 3B) (Tables S1 and S2).

**Figure 3.**
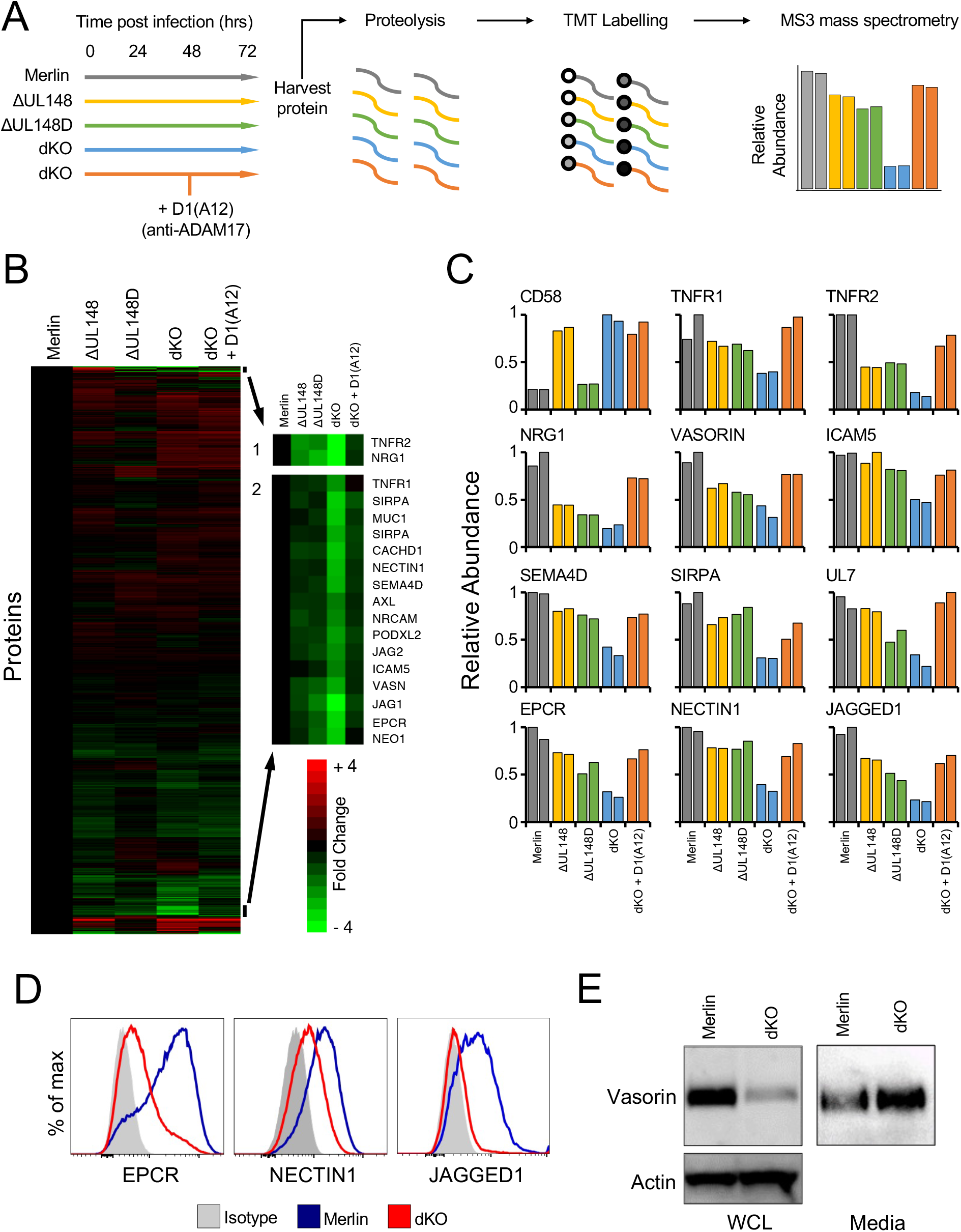
Legend: HCMV UL148 and UL148D upregulate numerous surface proteins by downregulating ADAM17. (A) Workflow of the PMP proteomics experiment on HF-TERT cells infected with the indicated HCMV strains for 72 h. A subset of the data from this experiment was used in Fig. 4A. The illustrated workflow shows the whole experiment. (B) Hierarchical cluster analysis of all proteins quantified in the experiment. (C) Examples of quantified proteins. CD58 is known to be targeted by UL148 in an ADAM17-independent manner. (D and E) Validation of PMP-identified proteins via flow cytometry (D) and immunoblotting (E) using HF-TERTs infected with HCMV strain Merlin or the dKO mutant for 72 h.

CD58 was exceptional in being upregulated on the cell surface in an UL148-dependent, UL148D-independent, ADAM17-independent manner (Fig. 3C) (36). The pattern of TNFR2 regulation agreed with our preceding findings (Fig. 1), with the dKO virus showing substantially less surface expression than either single gene deletion mutant. Using a significance score of *p*<0.05, 114 proteins (Table S1) were recovered on the surface of dKO-infected cells in an ADAM17-dependent fashion.

A number of these findings were validated orthogonally. Flow cytometry showed that EPCR and nectin1, highly expressed on human fibroblasts, were downregulated when UL148 and UL148D were deleted (Fig. 3D). Some of the very highly significant hits (p<0.00001; Table 1) were, however, not found at high levels on fibroblasts (data not shown). When these low-expressing proteins were over-expressed in fibroblasts by lentiviral transduction, they also showed the same pattern. Thus, downregulation of surface jagged1 on dKO-infected cells was readily detected by flow cytometry (Fig 3D), while Western blotting showed high levels of vasorin in Merlin-infected cells that were heavily reduced following infection with dKO HCMV. An inverse pattern was observed for vasorin in the supernatants from infected cells (Fig. 3E). Thus, while the modulation of TNFR2 led us to discover UL148 and UL148D’s control of ADAM17, their impact on the host cellular proteome was considerably more profound. Overall, impairment of ADAM17 expression by HCMV UL148 and UL148D resulted in global changes to the levels of numerous cell surface proteins with a concomitant inverse effect on their soluble forms.

**Table 1.**
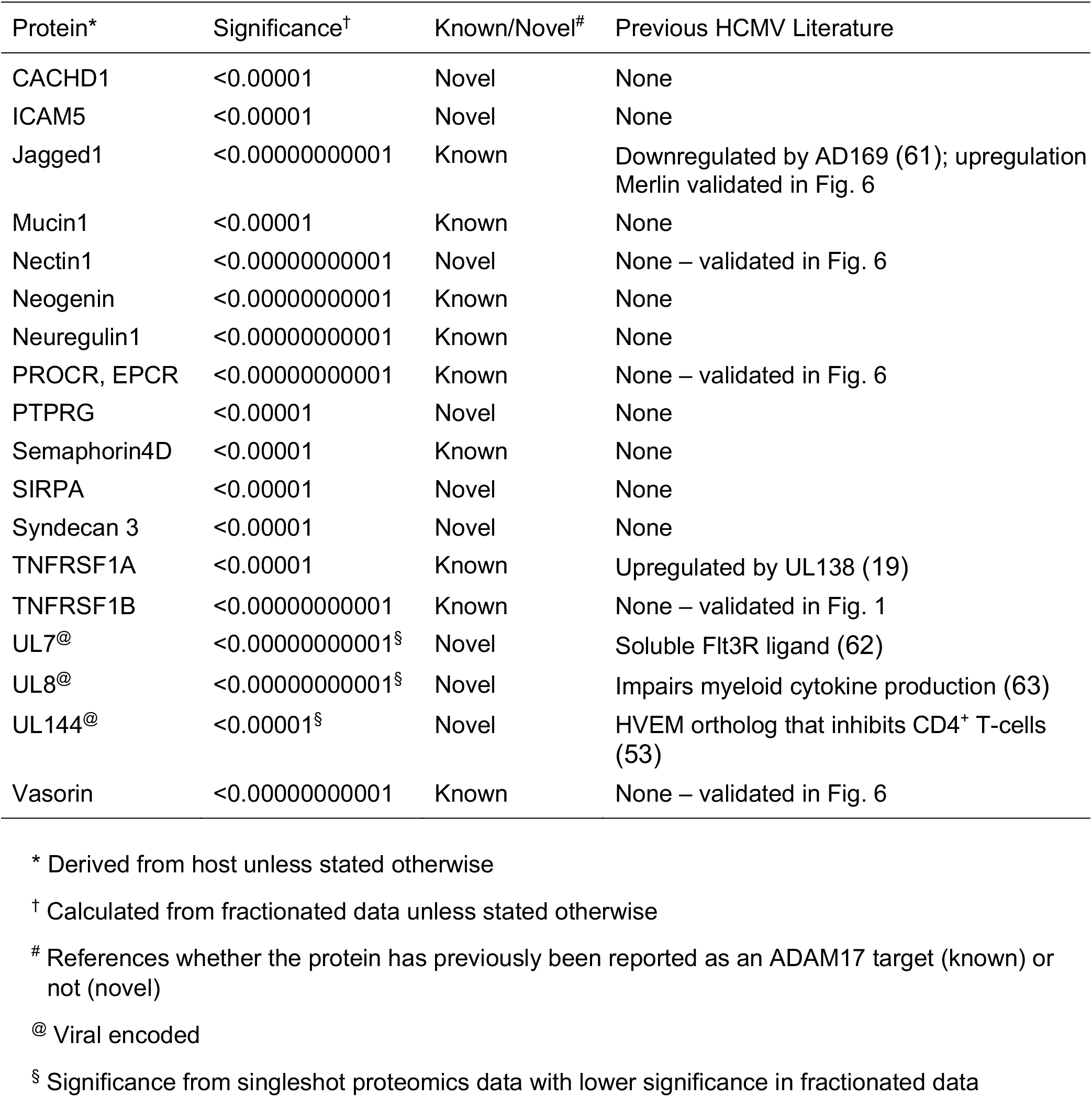
Summary of highly significant PMP protein hits (*p*<0.00001) stabilized on the surface of HCMV-infected cells through targeting of ADAM17

### ADAM17-dependent modulation of TNFα-induced responses during HCMV infection

We next investigated some of the functional consequences of these global alterations on HCMV-infected cells through blocking of ADAM17 via the application of D1(A12). Signalling through the TNFα/TNFR1/R2 axis, was measured using TNFα-induced cytokine production. Consistent with the lack of expression of ADAM17 and therefore the inability of D1(A12) to change TNFR2 expression on Merlin-infected cells (Fig. 2E), blocking of ADAM17 did not significant alter TNFα-induced cytokine production following infection with Merlin (Fig. 4A). In contrast, blocking of ADAM17 function on dKO-infected cells significantly raised levels of surface TNFR2 (Fig. 2E), and led to substantial increases in TNFα-induced cytokine production, with some such as IL-8 reaching over 40-fold higher than unstimulated levels (Fig. 4B). Thus, during HCMV infection, the inhibition of ADAM17 function promoted TNFα-signalling correlating with increased surface TNFR1 and 2.

**Figure 4.**
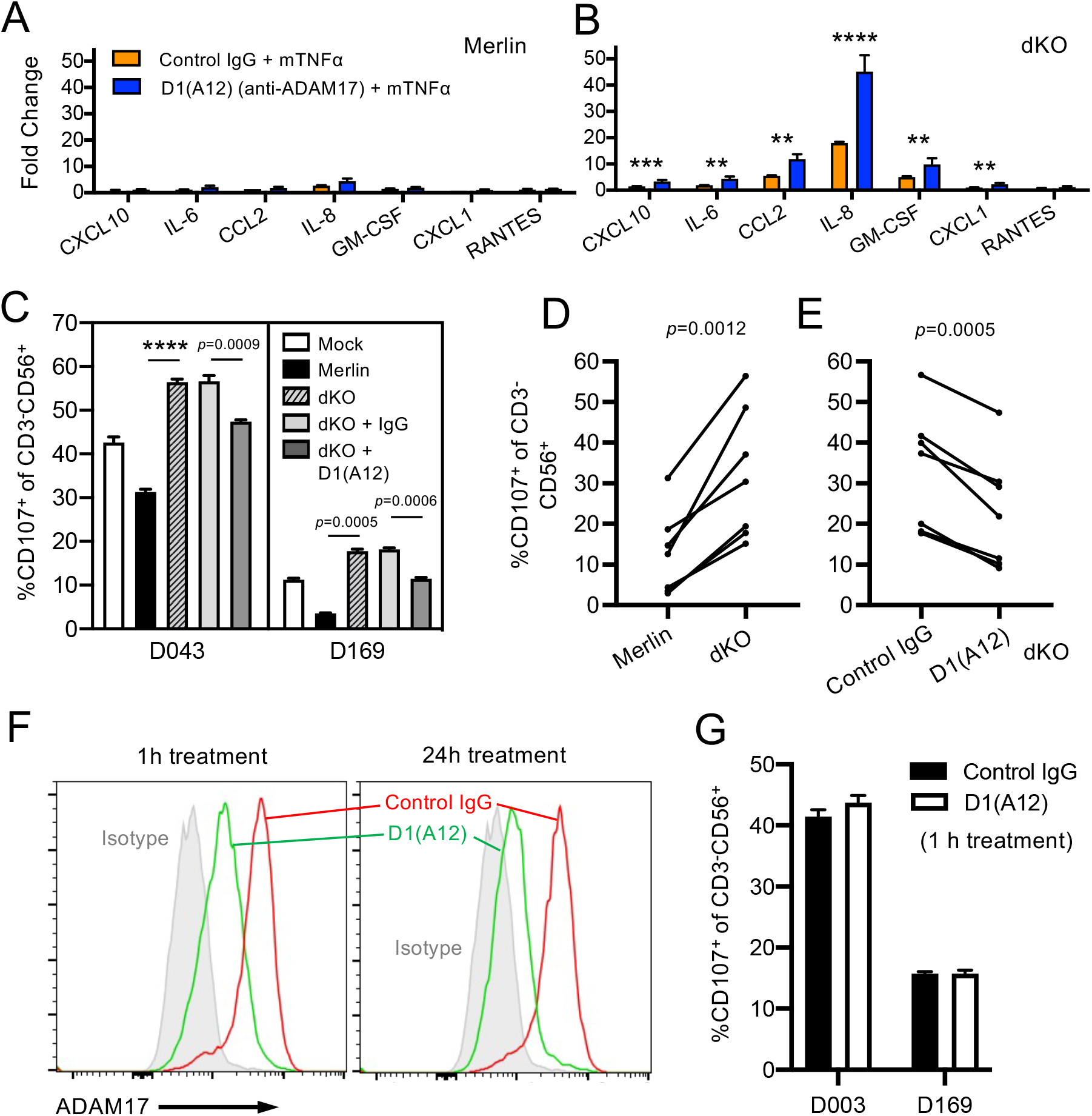
Legend HCMV UL148 and UL148D modulate multiple immune pathways in an ADAM17-dependent fashion. (A) HF-TERT cells were infected with HCMV strain Merlin or (B) the dKO mutant for 72h. Cells were treated with anti-ADAM17 antibody D1(A12) or control human IgG for 24 h prior, and TNFα for 18 h prior, to harvest of supernatants. Levels of IL-6, IL-8, GM-CSF, CXCL-10, CCL-2, CXCL-1, ICAM-1, VCAM, and RANTES were determined using bead arrays. Bars show mean ± SEM of triplicate infections. Two-way ANOVA with Tukey’s multiple comparison post-hoc test showed significance at \*\*\*\**p*<0.0001, \*\*\**p*<0.001, or \*\**p*<0.01. (C) Activation of NK lines from 2 different donors against HF-TERT cells, mock-infected or infected with the indicated HCMV strains. dKO-infected cells were further treated with D1(A12) or control human IgG 24 h prior to assay. Effector:target ratio of 10:1 used. Data are mean + SEM of quadruplicate samples. Brown-Forsythe ANOVA with Dunnett’s T3 multiple comparison post-test showed significance at \*\*\*\**p*<0.0001, or as indicated. Summary activation data from 7 NK lines against (D) HF-TERT cells infected with Merlin or dKO mutant, and (E) dKO-infected cells treated with control IgG or D1(A12). Each data point represents mean of quadruplicate samples. Paired *t*-Tests showed significance as indicated. (F) Flow cytometric histogram overlays showing ADAM17 expression after treatment with D1(A12) for 1 or 24 h. (G) NK activation of NK lines against dKO-infected cells after D1(A12) treatment for 1 h.

### ADAM17-dependent NK cell inhibition during HCMV infection

We further examined how these changes to the cell surface proteome altered interactions with effector immune cells important in immunity against HCMV. dKO infection resulted in significant increases in activation of NK cell lines from multiple donors compared to Merlin-infected cells (Fig. 4C, D), indicating that pathways targeted by UL148 and UL148D aid in evading the NK response. Application of D1(A12) to dKO-infected cells resulted in a significant decrease in the NK cell response of all NK cell lines tested (Fig. 4E), demonstrating that ADAM17 activity significantly contributed to the increase in NK activation induced by dKO-infected cells. NK inhibition was likely due to stabilization of one or more NK inhibitory ligand(s) rather than ADAM17 acting as an activating NK ligand itself, since ADAM17 surface expression was already reduced by a 1h treatment of dKO-infected cells with D1 (A12) (Fig. 4F), at a timepoint when NK inhibition did not occur (Fig. 4G). These data reveal a novel NK immune evasion strategy that may provide additional counter selection pressure against the potential antiviral effects of stabilising TNFR1/2 and highlight the simultaneous regulation of multiple immunological pathways caused by the targeting of ADAM17 during HCMV infection.

## DISCUSSION

In this study, we describe the profound effects that two genes, UL148 and UL148D, have on the cell surface proteome of, and soluble proteins produced by, HCMV-infected cells. UL148 and UL148D achieved this through their targeting of ADAM17, expression of a functionally active cell-surface version of which was markedly upregulated during HCMV infection in the absence of the 2 genes. A previous report describing a lack of ADAM17 modulation in HCMV-infected cells can be attributed to the use of strain AD169, which is missing the U_L_/b’ region that contains UL148 and UL148D (37).

Despite the similarity of their names, UL148 and UL148D are genetically unrelated and exhibit no overt homology to each other or any other HCMV gene (38). Both are ancient genes with homologs found in chimpanzee CMV (39). This work describes a first biological function for UL148D, while providing an additional role for UL148, which has been ascribed multiple functions previously. HCMV UL148 acts as an immune evasion gene via its intracellular retention of the cell adhesion molecule CD58 (36), but also activates the unfolded protein response involved in ER stress signalling, binding a key regulator of the ER-dependent degradation pathway (ERAD) Suppressor/Enhancer of Lin-12-like protein (Sel1L), thereby altering degradation of the viral envelope glycoprotein gO and changing viral tropism ((31, 32, 40)). Neither intracellular retention, nor triggering of the ERAD pathway, seem to be involved in the targeting of ADAM17 by UL148 or UL148D, both of which may use distinct mechanisms. The proteins have distinct expression profiles, with UL148D being expressed earlier than UL148 during infection and displaying temporal protein profile Tp2 kinetics, whereas UL148 is predominantly expressed later, with temporal protein profile Tp5 kinetics (41). Although both proteins have ER-retention motifs (RRR at residues 314-316 for UL148 and IRR at residues 27-29 for UL148D) (elm.eu.org), they did not co-localize within infected cells and interactome studies have not found any direct interaction between ADAM17 and UL148 or UL148D (42). Our data also showed that ADAM17 expression was not rescued during wildtype HCMV infection by MG132, a proteasomal inhibitor that acts downstream of the ERAD pathway. ADAM17 expression and function is, however, complex, involving close regulation by multiple chaperones (iRhom1, iRhom2, FRMD8) and a Furin-dependent cleavage event. It remains to be seen whether UL148 and UL148D may indirectly disrupt ADAM17 processing through interactions with these complexes or possibly other undiscovered chaperones.

The primary significance of this study is the characterization of the broad regulation of the cell surface proteome and its simultaneous impact on multiple biological responses induced by impairment of ADAM17. Its targeting by 2 distinct HCMV genes suggests countering this pathway is important for HCMV biology, while ADAM17’s numerous substrates highlight its potential to act as a novel regulatory hub. In line with this, virus-mediated downregulation of ADAM17 altered expression of dozens of proteins with our study identifying many new ADAM17 substrates. Exploring the 18 very highly significant (*p*<0.00001) proteins summarized in Table 1, nine were known (26, 43), while the remaining nine were novel, including nectin1, PTPRG and SIRPA, which have previously been reported as targeted by ADAM10 (44–46). Using *p*<0.05 as a cut-off, the total number of ADAM17 substrates stabilized by wildtype HCMV infection numbered 114, one hundred and one of which have not previously been reported to be cleaved by ADAM17 (Table S1). Some known substrates failed to show ADAM17-dependent modulation. This may reflect cell-type and context-dependent ADAM17 shedding activity (47), however, HCMV also counteracts the stabilization of specific ADAM17 substrates through independent mechanisms. For example, MICA and MICB are ligands for the activating receptor NKG2D, and their ADAM17-dependent stabilization would render cells susceptible to NK cell attack. Therefore, MICA is targeted for lysosomal degradation via US18 and US20 (35) and proteasomal degradation via UL147A (48). HCMV UL142 and UL16 additionally retain MICA and MICB, respectively, in the cis-Golgi (49, 50). These observations suggest that the upregulation of certain host proteins that would occur due to viral inhibition of ADAM17 are blocked via additional functions if unfavourable for HCMV. Likely of direct virological significance, a number of viral proteins were also stabilized on the surface of HCMV-infected cells following targeting of ADAM17, including UL7, UL8, and UL144 in the p<0.0001 significance group, while increasing to 9 when using p<0.05 as a cut-off. These viral proteins may require surface expression for their intended functions.

The impairment of ADAM17 expression impacted at least two anti-viral immune pathways, resulting in increased TNFα-induced cytokine responses but reduced NK cell activation. The underlying mechanism for ADAM17-dependent NK inhibition is distinct from the previously described NK and T-cell inhibitory function of UL148 mediated via the intracellular retention of CD58 (36), because CD58 expression was not dependent on ADAM17. Indeed, several of the highly significant hits for ADAM17-dependent stabilization during HCMV infection have previously been reported as NK inhibitors, such as MUC1 (51) and nectin1 (52). Furthermore, multiple viral protein hits also have established or potential NK inhibitory activity; B and T lymphocyte attenuator (BTLA) is the ligand for UL144 (53) and has recently been implicated in NK cell immunosuppression (54), while UL40 (*p*<0.0002; Table S1), is a recognized inhibitor of NKG2A^+^ NK cells (55, 56).

The functional consequences of ADAM17 dysregulation, however, are likely to go beyond NK cell inhibition. For example, both jagged1 and vasorin play roles in the development of Tregs (57–59), and both were upregulated on the surface of Merlin-infected cells. Treg cells are immunosuppressive and impair protective immunity in a number of viral infections, including herpes simplex virus, HIV, and hepatitis C virus (60). A similar scenario may therefore apply to HCMV. A previous report suggesting that HCMV downregulated jagged1 (61) may be explained by the use of HCMV strain Towne, which like strain AD169, lacks UL148 and UL148D (38). Furthermore, all 3 viral proteins showing highly significant cell surface stabilization due to UL148/UL148D have immune functions, with UL7 promoting myelopoiesis as a ligand for Fms-like tyrosine kinase 3 receptor (62), UL8 impairing myeloid proinflammatory cytokine production (63), and UL144 inhibiting CD4^+^ T-cell proliferation through its interaction with BTLA (53).

The significance of the extensive modulation of the infected cell surface is likely to manifest beyond immunoregulation. Although ADAM17 is expressed to the greatest extent in cells of the lymphatic system, it is also found in most other tissues and is developmentally regulated from conception to death (64). Indeed, ADAM17 deficiency is lethal in embryonic mice (65). Furthermore, neuroprotective properties have been attributed to ADAM17, alongside roles in repair of the brain and central nervous system (CNS) (25). It is tempting to speculate that HCMV’s targeting of ADAM17 may contribute to the neurodevelopmental abnormalities associated with congenital CMV (cCMV) infection, with ~90% of symptomatic neonates suffering from damage caused to the CNS resulting in mental retardation, vision and hearing loss (66, 67). Further research is needed to explore this concept and whether ADAM17 activation may represent a stress-induced cellular response to infection considering the high levels of intracellular ADAM17 (68), its activation at the cell surface following infection and the rapidity that HCMV targets its expression with downregulation even within 6 hours of infection (41). As such, it may be a common target for modulation by diverse microorganisms as *Lactobacillus gasseri*, has also been shown to suppress pro-inflammatory cytokine production through inhibiting ADAM17 expression (69).

## Materials and methods

### Cell lines

Human fetal foreskin fibroblasts were immortalised with human telomerase (HF-TERT), HF-TERT cells transfected with the Coxsackie-adenovirus receptor (HF-CAR) were described previously(70). Cells were maintained in Dulbecco’s minimal essential medium (DMEM) supplemented with 10% fetal calf serum (FCS) at 37 °C/5% CO2.

### Viruses and viral infection

HCMV strain Merlin RCMV1111/KM192298 (RL13-, UL128-) and Merlin recombinants containing single gene deletions in UL/b’ were generated as described previously(71). HCMV containing C-terminal epitope tags on UL148 and UL148D were generated as described previously and verified by next generation sequencing after reconstitution(35, 72). UL148-V5-tagged HCMV (RCMV2445) was made previously(36) and used to HA-tag UL148D, generating a double-tagged HCMV (RCMV2929). En passant mutagenesis method was used to tag UL148D as described previously(73), using UL148DF (TTTACGCAGCAGCAGGCACGCAACGGGAGCGGCAGCGGCAGCGCTTACCCCTACGACGTGCCCGACTACGCCTAGACAATAGGGATAACAGGGTAATGGC) and UL148D R (CCGGCTACGGCGCTTGGAGCTGTAGCCGCCTGGGACTTGTCTAGGCGTAGTCGGGCACGTCGTAGGGGTAAGCGCTTCAGAAGAACTCGTCAAGAAGGCG) primers. Oligonucleotide primers were purchased from Eurofins. Recombinant adenovirus vectors (RAds) expressing individual HCMV UL148 and UL148D genes were generated as described previously (74). Viral infections were performed as described previously (35). For degradation inhibition studies, inhibitors of the proteasomal (MG132, 10 μM, Merck) and lysosomal degradation pathways (leupeptin, 200 μM, Merck) were added to the cells 12 h prior to harvest. For ADAM17 blocking studies, anti-ADAM17 (clone D1(A12), Abcam) or human IgG isotype (polyclonal, Abcam) were added at a final concentration of 100 nM for 24 h and washed away prior to harvest.

### Immunofluorescence

Cells were seeded and infected in glass-bottom 96-well plates (Ibidi) as described previously (71). Samples were fixed with 4% paraformaldehyde and permeabilized with 0.5% NP-40. Primary antibodies included anti-V5 tag (clone SV5-Pk1, Abcam) and anti-HA tag (clone 2-2.2.14, Thermo Fisher Scientific). Secondary antibodies included Alexa Fluor 488-conjugated anti-rabbit IgG (polyclonal, Thermo Fisher Scientific) and Alexa Fluor 594-conjugated anti-mouse IgG (polyclonal, Thermo Fisher Scientific). Nuclear staining was performed using Hoechst.

### Flow Cytometry

Adherent cells were harvested with TrypLE Express (Thermo Fisher Scientific), stained with the relevant antibodies, fixed with 4% paraformaldehyde, and analyzed on an Accuri C6 flow cytometer (BD Biosciences) or an Attune NxT flow cytometer (Thermo Fisher Scientific). Antibodies and reagents used for flow cytometry staining included Live/Dead Aqua (Thermo Fisher Scientific), Live/Dead eFluor 660 (Thermo Fisher Scientific), anti-CD120b (clone REA520, Miltenyi Biotec), anti-EPCR (clone RCR-401, BioLegend), anti-nectin1 (clone CK41, BD Biosciences), anti-CD3 (clone HIT31, BioLegend), anti-CD107a (clone H4A3, BioLegend), anti-CD56 (clone 5.1H11, BioLegend), anti-CD8a (clone RPA-T8, BioLegend), anti-Jagged1 (clone 4A24, GeneTex), anti-ADAM17 (clone 111633, R&D Systems), anti-mouse IgG-Alexa Fluor 647 (polyclonal, Thermo Fisher Scientific), and anti-rabbit IgG-Alexa Fluor 647 (polyclonal, Thermo Fisher Scientific). Apoptosis assays were perfomed using CellEvent Caspase 3/7 Green Detection Reagent (Thermo Fisher Scientific) and Live/Dead eFluor 660 (Thermo Fisher Scientific). Cells were treated with TNFα at a final concentration of 30 ng/ml 48 h prior to harvest. Data were analyzed using Accuri C6, Attune NxT, or FlowJo V10 softwares. Cytokine release was quantified using a LEGENDplex Assay (BioLegend). Cells were treated with TNFα at a final concentration of 30 ng/ml 18 h prior to supernatant collection. Beads were analyzed by flow cytometry using LEGENDplex data analysis software (BioLegend).

### Immunoblotting

Protein lysates were prepared and separated as described previously(36). Primary antibodies included anti-V5 tag (rabbit polyclonal, Abcam) and anti-HA tag (clone 2-2.2.14, Thermo Fisher Scientific), anti-ADAM17 (rabbit polyclonal, Abcam), anti-actin (rabbit polyclonal, Sigma), anti-CD120b (clone EPR1653, abcam), anti-Vasorin (clone 4G7, Novus Biologicals), anti-MICA/B (clone BAM01), and anti-nectin2 (clone EPR6717, Abcam). Secondary antibodies included anti-mouse IgG-HRP (polyclonal, Bio-Rad) and anti-rabbit IgG-HRP (polyclonal, Bio-Rad). For EndoH and PNGaseF digestion, enzymes (New England Biolabs) were incubated with cell lysates for 18 h at 37°C prior to protein reduction. For ADAM17 immunoblotting, lysates were incubated with concanavalin A beads for 3 h at 4°C, followed by elution of glycosylated ADAM17 forms and protein reduction.

### Plasma membrane profiling

Plasma membrane profiling was performed as described previously with minor modifications, to cells infected in biological duplicate(41, 75). After washing cells infected in duplicate, surface sialic acid residues were oxidized with sodium-meta-periodate and labelled with aminooxy-biotin, and after quenching the reaction, biotinylated cells were scraped into 1% (v/v) Triton X-100. Biotinylated glycoproteins were enriched with high-affinity streptavidin agarose beads and washed extensively. Captured protein was denatured with DTT, alkylated with iodoacetamide (IAA), and digested on-bead with trypsin for 3 h in 100 mM HEPES pH 8.5. Tryptic peptides were collected and subjected to labelling with tandem mass tags. The following labels were applied: 126 wild-type #1, 127N wild-type #2, 127C ΔUL148 #1, 128N ΔUL148 #2, 128C ΔUL148D #1, 129N ΔUL148D #2, 129C ΔUL148/ΔUL148D #1, 130N ΔUL148/ΔUL148D #2, 130C ΔUL148/ΔUL148D+D1 (A12) #1, and 131N ΔUL148/ΔUL148D+D1(A12) #2. Labelled peptides were combined in a 1:1:1:1:1:1:1:1:1:1 ratio, enriched, and then subjected to high-pH reversed-phase fractionation. Fractions were combined for analysis by recombining all wells from sets of two adjacent columns in the resulting 96-well plate (*i.e*., combining wells in columns A+B, C+D, E+F etc). This resulted in six fractions and an unfractionated ‘singleshot’ sample for analysis via mass spectrometry (MS).

### LC-MS3

Mass spectrometry data were acquired using an Orbitrap Fusion Lumos (Thermo Fisher Scientific) with an UltiMate 3000 RSLC nano UHPLC equipped with a 300 μm ID × 5 mm Acclaim PepMap μ-Precolumn (Thermo Fisher Scientific) and a 75 μm ID × 50 cm 2.1 μm particle Acclaim PepMap RSLC Analytical Column (Thermo Fisher Scientific). Loading solvent was 0.1% trifluoroacetic acid (TFA), analytical solvent A was 0.1% formic acid (FA), and analytical solvent B was acetonitrile (MeCN) + 0.1% FA. All separations were carried out at 55°C. Samples were loaded at 10 μl/minute for 5 minutes in loading solvent before beginning the analytical gradient. All samples were run with a gradient of 3-34% B, followed by a 5 minute wash at 80% B, a 5 minute wash at 90% B and equilibration for 5 minutes at 3% B. Each analysis used a MultiNotch MS3-based TMT method (76). The following settings were used: MS1: 400–1400 Th, quadrupole isolation, 120,000 resolution, 2 x 10^5^ automatic gain control (AGC) target, 50 ms maximum injection time, and ions injected for all parallizable time; MS2: quadrupole isolation at an isolation width of m/z 0.7, collision-induced dissociation (CID) fragmentation with normalized collision energy (NCE) 30 and ion trap scanning out in rapid mode from m/z 120, 1×104 AGC target, 70 ms maximum injection time, and ions accumulated for all parallizable time in centroid mode; MS3: in synchronous precursor selection mode, the top 10 MS2 ions were selected for higher energy collisional dissociation (HCD) fragmentation (NCE 65) and scanned in the Orbitrap at 50,000 resolution with an AGC target of 5 × 10^4^ and a maximum accumulation time of 150 ms, and ions were not accumulated for all parallelizable time. The entire MS/MS/MS cycle had a target time of 3 s. Dynamic exclusion was set to ± 10 ppm for 90 s. MS2 fragmentation was trigged on precursors at 5 x 10^3^ counts and above.

### Data analysis

Mass spectra were processed using a Sequest-based software pipeline for quantitative proteomics, “MassPike”, through a collaborative arrangement with Professor Steven Gygi’s laboratory at Harvard Medical School. Spectra were converted to mzXML using an extractor built upon Thermo Fisher Scientific’s RAW File Reader Library (version 4.0.26). In this extractor, the standard mzxml format has been augmented with additional custom fields that are specific to ion trap and Orbitrap MS data and essential for TMT quantitation. These additional fields include ion injection times for each scan, Fourier transform-derived baseline and noise values calculated for every Orbitrap scan, isolation widths for each scan type, scan event numbers, and elapsed scan times. This software is a component of the MassPike software platform licensed by Harvard Medical School.

A combined database was constructed from (a) the human Uniprot database (26 January 2017), (b) the HCMV strain Merlin Uniprot database, (c) all additional non-canonical HCMV(77), (d) a six-frame translation of HCMV strain Merlin filtered to include all potential ORFs of ≥8 amino acids (delimited by stop-stop rather than requiring ATG-stop) and (e) common contaminants such as porcine trypsin and endoproteinase LysC. ORFs from the six-frame translation (6FT-ORFs) were named as follows: 6FT_Frame_ORFnumber_length, where Frame is numbered 1-6, and length is the length in amino acids. The combined database was concatenated with a reverse database composed of all protein sequences in reversed order. Searches were performed using a 20 ppm precursor ion tolerance. Fragment ion tolerance was set to 1.0 Th. TMT tags on lysine residues and peptide N termini (229.162932 Da) and carbamidomethylation of cysteine residues (57.02146 Da) were set as static modifications, while oxidation of methionine residues (15.99492 Da) was set as a variable modification.

A target-decoy strategy was employed to control the fraction of erroneous protein identifications(78). Peptide spectral matches (PSMs) were filtered to an initial peptide-level false discovery rate (FDR) of 1% with subsequent filtering to attain a final protein-level FDR of 1 %. PSM filtering was performed using a linear discriminant analysis, as described previously(78). This distinguishes correct from incorrect peptide identifications (IDs) in a manner analogous to the widely used Percolator algorithm (https://noble.gs.washington.edu/proj/percolator/) by employing a distinct machine learning algorithm. The following parameters were considered: XCorr, ΔCn, missed cleavages, peptide length, charge state, and precursor mass accuracy.

Protein assembly was guided by the principles of parsimony to produce the smallest set of proteins necessary to account for all observed peptides (algorithm described in(78)). Where all PSMs from a given HCMV protein could be explained either by a canonical gene or non-canonical ORF, the canonical gene was picked in preference.

In a few cases, PSMs assigned to a non-canonical gene or 6FT-ORF were a mixture of peptides from the canonical protein and the ORF. In these cases, the peptides corresponding to the canonical protein were separated from those unique to the ORF, generating two separate entries.

Proteins were quantified by summing TMT reporter ion counts across all matching peptide-spectral matches using “MassPike”, as described previously(76). Briefly, a 0.003 Th window around the theoretical m/z of each reporter ion (126, 127n, 127c, 128n, 128c, 129n, 129c, 130n, 130c, and 131n) was scanned for ions, selecting the maximum intensity nearest to the theoretical m/z. The primary determinant of quantitation quality is the number of TMT reporter ions detected in each MS3 spectrum, which is directly proportional to the signal-to-noise (S:N) ratio observed for each ion. Conservatively, every individual peptide used for quantitation was required to contribute sufficient TMT reporter ions (minimum of ~1250 per spectrum), so that each on its own could be expected to provide a representative picture of relative protein abundance(76). An isolation specificity filter with a cutoff of 50% was additionally employed to minimise peptide co-isolation(76). Peptide-spectral matches with poor quality MS3 spectra (>9 TMT channels missing and/or a combined S:N ratio of <250 across all TMT reporter ions) or no MS3 spectra were excluded from quantiration. Peptides meeting the stated criteria for reliable quantitation were then summed by parent protein, in effect weighting the contributions of individual peptides to the total protein signal based on their individual TMT reporter ion yields. Protein quantiration values were exported for further analysis in Excel.

For protein quantitation, reverse and contaminant proteins were removed, and each reporter ion channel was summed across all quantified proteins and normalized on the assumption of equal protein loading across all channels. Fractional TMT signals, reporting the fraction of the maximal signal observed for each protein in each TMT channel rather than the absolute normalized signal intensity, were used for further analysis and visualization. This approach effectively corrected for differences in the numbers of peptides observed per protein. Normalized S:N values are presented in Tables S1 and S2, assuming equal protein loading across all samples. As it was not possible to assign peptides to HLA-A, HLA-B, or HLA-C alleles with confidence, S:N values were further summed to give a single combined result for HLA-A, HLA-B or HLA-C.

Hierarchical centroid clustering was based on an uncentered Pearson correlation and visualised using Java Treeview (http://jtreeview.sourceforge.net). p-values for protein fold change were estimated using the method of Significance B, calculated in MaxQuant and corrected for multiple hypothesis testing using the method of Benjamini-Hochberg(79).

### Soluble TNFR2 detection

Soluble TNFR2 levels were measured using a Human TNFR2 Quantikine ELISA (R&D Systems). Optical density was measured using a FLUOstar Omega Microplate Reader (BMG LABTECH).

### NK cell lines and CD107a degranulation assays

CD14^-^CD3^-^CD56^+^ NK cells were purified directly *ex vivo* via FACS and stimulated with γ-irradiated allogeneic PBMCs and LCL-721.221 cells (1:1 ratio) and PHA-P (10 μg/ml) in RPMI-1640 medium supplemented with 10% FCS, 5% human AB serum (Welsh Blood Service), 100 U/ml penicillin, 0.1 mg/ml streptomycin, 2mM L-glutamine, 100 U/ml rhIL-2, and 10 ng/ml IL-15 (NK cell medium) for 3 days at 37 °C. Lines were maintained at 1–2 × 10^6^ cells/ml by replenishing NK cell medium every 3–4 days. Rested cell lines were harvested for functional assays after 2 weeks in culture. The purity of all cell lines was >96%. NK cell degranulation assays were performed as described previously(36, 80), using an effector:target ratio of 10:1. Flow cytometry analysis was used to identify responding CD3^-^CD56^+^ NK cells.

### Statistical analysis

Statistical significance was determined using one-or two-way ANOVAs, with Bonferroni, Tukey’s or Dunnett’s T3 multiple comparison post-hoc tests as appropriate. NK CD107a degranulation summary data was analyzed using a paired *t*-Test after data had passed Shapiro-Wilk normality testing. Statistical analysis was performed using GraphPad Prism software. *P*-values of <0.05 were considered significant.

## Data and materials availability statement

The proteomics data have been uploaded to the ProteomeXchange Consortium (http://www.proteomexchange.org/) via the PRIDE(81) partner repository. All materials described in this manuscript and full protocols can be obtained on request from the corresponding author.

## Ethics statement

Healthy adult volunteers provided blood for this study after giving written informed consent in accordance with the principles of the Declaration of Helsinki. The study was approved by the Cardiff University School of Medicine Research Ethics Committee (reference numbers 10/20 and 16/52).

## Acknowledgements

We are grateful to Prof. Steve Gygi for providing access to the “MassPike” software pipeline for quantitative proteomics. This work was funded by the Medical Research Council (MR/P001602/1 to E.C.Y.W., P.T., and G.W.G.W.; MR/V000489/1 to E.C.Y.W., D.A.P., R.J.S., and S.K.; MR/S00971X/1 to R.J.S. and E.C.Y.W.; MC_UU_12014/3 to A.J.D.) and further supported by two Cardiff University PhD Studentships (one part-funded by the Medical Research Council and one from the Systems Immunity University Research Institute). D.A.P. was supported by a Wellcome Trust Senior Investigator Award (100326/Z/12/Z). M.P.W. was supported by a Wellcome Trust Senior Clinical Research Fellowship (108070/Z/15/Z). An Attune flow cytometer (ThermoFisher) was used throughout which was obtained and serviced with the following grants - MR/P001602/1, MR/S00971X/1, MR/V000489/1, and Wellcome Trust grants 204870 (awarded to P Griffiths, UCL) and 207503/Z/17/Z (awarded to I Humphreys, Cardiff University). For the purpose of open access, the author has applied a CC BY public copyright licence to any Author Accepted Manuscript version arising from this submission.

## Author Contributions

Conceptualization: E.C.Y.W., A.R., M.Patel

Data curation: E.C.Y.W., R.J.S., M.P.W.

Formal Analysis: A.R., M.Patel., E.C.Y.W., K.N., M.P.W.

Funding acquisition: E.C.Y.W., G.W.G.W., P.T., R.J.S., D.P., S.K., A.J.D., M.P.W.

Investigation: A.R., M.Patel., K.N., M.Potts., C.A.F., S.K., B.L., J.N., K.L.M., K.L., L.N., T.M.T., J.P.T., V.M.V., D.R.

Methodology: E.C.Y.W., R.J.S., M.P.W., S.K., A.J.D., C.A.F., B.L., K.L.M., K.L., J.N., D.R. Project administration: E.C.Y.W., D.R.

Resources: K.L.M., K.L., D.A.P., A.J.D.

Supervision: E.C.Y.W, R.J.S., G.W.G.W.

Validation: J.N., A.J.D.

Visualization: A.R., M.P., E.C.Y.W., K.N.

Writing – original draft: A.R., E.C.Y.W.

Writing – review & editing: A.R., E.C.Y.W., D.A.P., R.J.S., M.P.W., C.A.F., G.W.G.W.

## Notes

### Competing Interest Statement

The authors have declared no competing interest.

