## Supplemental Figure 1 for "ADAM17 targeting by human cytomegalovirus remodels the cell surface proteome to simultaneously regulate multiple immune pathways"

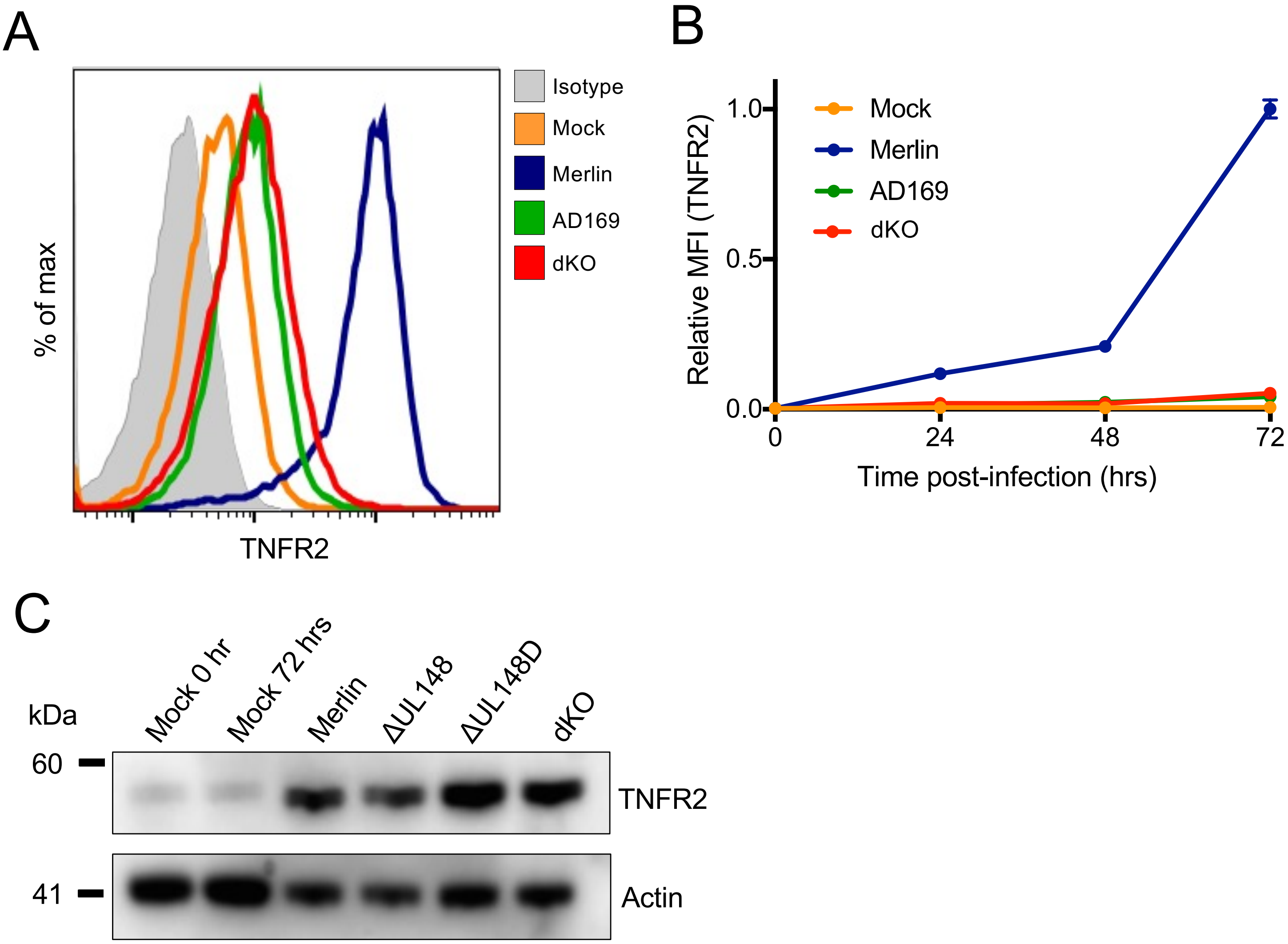


**Supplemental Figure 1. UL148 and UL148D act together to upregulate surface expression of TNFR2.** HF-TERT cells were mock-infected or infected with the indicated HCMV strains and analyzed by flow cytometry at 72 h pi (A) or at indicated time points pi (B) for surface expression of TNFR2. MFI values are shown relative to Merlin-infected cells (set to 1) at 72 h pi. (C) HF-TERT cells were mock-infected or infected with HCMV strain Merlin or the indicated deletion mutants, and whole-cell lysates were analyzed by immunoblotting at 72 h pi for TNFR2. Actin was used as a loading control.
